# Diagnostics of dated phylogenies in microbial population genetics

**DOI:** 10.1101/2025.09.24.678031

**Authors:** Xavier Didelot, Jake Carson, Paolo Ribeca, Erik Volz

**Affiliations:** School of Life Sciences, University of Warwick, United Kingdom; Department of Statistics, University of Warwick, United Kingdom; Mathematics Institute, University of Warwick, United Kingdom; UK Health Security Agency, London, United Kingdom; Biomathematics and Statistics Scotland, The James Hutton Institute, Edinburgh, United Kingdom; MRC Centre for Global Infectious Disease Analysis, Department of Infectious Disease Epidemiology, Imperial College London, London, United Kingdom

## Abstract

Microbial population genetic studies often involve the use of a dated phylogeny to show how the genomes are related over a relevant timescale. Many tools have recently been developed to date the nodes of a standard phylogeny, but all make underlying assumptions that may not be realistic for a given dataset, making the results potentially unreliable. Model comparison is sometimes used to remedy this issue, whereby inference under several models is compared to establish which result can be trusted. Although such comparison is clearly useful to assess the relative merits of several inference attempts, here instead we focus on the problem of evaluating how good an inference is in absolute terms, without comparison. We consider several approaches for diagnosing potential issues in a reconstructed dated phylogeny, including outlier detection, posterior predictive checking and residual analysis. These methods are well-established diagnostics tools in other areas of statistics, but here we show how they can be applied to the specific inference of dated phylogenies. We illustrate their use on many simulated datasets, with inference being performed either from the correct model to quantify the specificity or from an incorrect model to quantify the sensitivity of the diagnostics methods. We also applied the methods to three real-life datasets to showcase the range of issues that they can detect. We advocate the use of these diagnostics tools for all microbial population genetic studies that involve the reconstruction of a dated phylogeny.

## INTRODUCTION

Dated phylogenies, also known as tip-calibrated, time-stamped or time-calibrated phylogenies, have become a ubiquitous tool in the study of microbial population genetics (Drummond et al. 2003; Biek et al. 2015; Rieux and Balloux 2016). In a dated phylogeny, the branch lengths are measured in a unit of time, for example years or days, rather than a unit of evolution as in a standard phylogeny. Consequently, the tips of a dated phylogeny are aligned with the (typically known) dates of sampled genomes and the internal nodes are aligned with the (typically inferred) dates of the last common ancestors between subsets of the genomes. Many tools exist to build dated phylogenies, either from a sequence alignment using for example BEAST ( Suchard et al.2018) or BEAST2 (Bouckaert et al. 2019), or by dating the nodes of a standard phylogeny, using for example LSD (To et al. 2016), node.dating (Jones and Poon 2017), treedater (Volz and Frost 2017), BactDating ( Didelot et al. 2018) or TreeTime ( Sagulenko et al.2018). In this paper we focus on the latter, that is dated phylogenies that have been estimated from an undated phylogeny and given the sampling dates of the leaves. The dated phylogeny is interesting in itself, since it depicts the ancestral relationships of sampled genomes over time and can provide an estimate of the dates of emergence of clades of interest. But dated phylogenies are also often used as the starting point for further analysis (Didelot and Parkhill 2022), such as inference of demographic history ( Volz and Didelot 2018), phylogeography (Roberts et al. 2025) or transmission between hosts ( Didelot et al.2017a).

There are many factors that can invalidate the results of a dated phylogenetics analysis. One potential issue that has been well studied is whether the temporal signal is strong enough for dating to be performed, or in other words whether the population is measurably evolving (Drummond et al. 2003; Biek et al. 2015). Several techniques have been proposed to evaluate the significance of the temporal signal, such as testing the correlation between sample dates and root-to-tip distances in an undated tree (Rambaut et al. 2016), comparing results with and without randomizing the sample dates (Duchene et al. 2015) and comparing results with correct sample dates and with all sample dates forced equal (Rambaut 2000; Duchene et al. 2020). Another possible source of problem concerns the confounding effect that population structure can have on dating ( Duchene et al. 2015; Murray 2016). This is especially true when the substructures are imbalanced (Duchêne et al. 2015), are sampled at different dates (Tong et al. 2018), have different clock rates (Wertheim et al. 2012) and when the population structure is strong ( Navascués and Emerson 2009). More generally, any dating method makes assumptions, sometimes unstated, and if these assumptions are not met the results can be incorrect. This includes the choice of a molecular clock model, which represents how mutations accumulate during the evolution of the population (Kumar 2005; Lepage et al. 2007). For Bayesian methods of dating there is also the need to specify an ancestral prior model, for example the heterochronous coalescent model with constant population size ( Drummond et al. 2002) or a birth-death process (Gernhard 2008).

One approach that has been used to ensure that there are no incorrect assumptions being made is to perform inference under multiple models and perform model comparison, typically by computing a Bayes Factor or an approximation of it (Baele et al. 2012; Li and Drummond 2012; Bouckaert and Drummond 2017). However, this requires multiple runs under different models, and only provides a relative measure of model appropriateness, with no indication of how good the best model actually is in absolute terms. In other words, it may be the case that inference under a given model is less incorrect than other attempts, but still not correct enough to be of any biological value.

Here we investigate an alternative approach, in which we seek to evaluate the correctness of an inference and detect if there are any reasons to believe that the inference is not valid. This approach is sometimes referred to as model checking, model criticism, model diagnostics or model validation, and it is complementary with the model comparison methodology mentioned above ( Carota et al. 1996). We investigate several methods for diagnosing potential problems in dated phylogenies, including outlier detection analysis, posterior predictive analysis and residual analysis. We use simulated datasets to test their ability to detect a range of problems in the inference, such as an incorrectly specified molecular clock model (Kumar 2005; Lepage et al. 2007) and the aforementioned confounding effect of population structure ( Murray et al. 2016). We also demonstrate that model criticism can be useful in practice to detect a wide range of issues that arise when dating real datasets from recent microbiology population genetics studies.

## RESULTS

### Outlier detection analysis

Before dating a phylogeny, it is useful to test the temporal signal by computing a linear regression between the known isolation dates of the tips and the distances from each tip to the root (assuming for now that the root is known). A frequently used software for this analysis is TempEst, which also allows the detection of outliers as potential problems (Rambaut et al. 2016). For example, in an analysis of 260 genomes from the current pandemic of *Vibrio cholerae*, 17 genomes were found to be outliers in the root-to-tip analysis which was explained by the fact they were hypermutators (Didelot et al. 2015). A similar outlier detection approach was implemented in treedater with the difference that it is applied to the distribution of likelihood per branch and compared to its expected distribution (Volz and Frost2017).

To illustrate the use of these outlier detection methods, a dated phylogeny was simulated including 100 leaves uniformly distributed between 2010 and 2020, under the heterochronous coalescent model Drummond et al. (2002) with constant population size *N*_e_*g* = 1 year. We applied a strict clock model (Equation 1) to this dated phylogeny, with clock rate *µ* = 10 substitutions per year, except that for five randomly selected leaves we added 20 substitutions, resulting in the phylogeny shown in Figure 1A. The analysis of root-to-tip distances is shown in Figure 1B, with all five modified tips clearly visible but close to the upper bound of the 95% expected envelope. When treedater was applied to this dataset, all five outliers were detected and given multiple testing corrected p-values between 8.1 *·* 10^−8^ and 3.9 *·* 10^−3^. Figure 1C shows for each branch of the treedater reconstructed tree its duration, number of substitutions and likelihood. The five outliers are clearly visible. When the five outliers were removed and treedater rerun, the resulting Figure 1D looked much more satisfactory.

**Figure 1:**
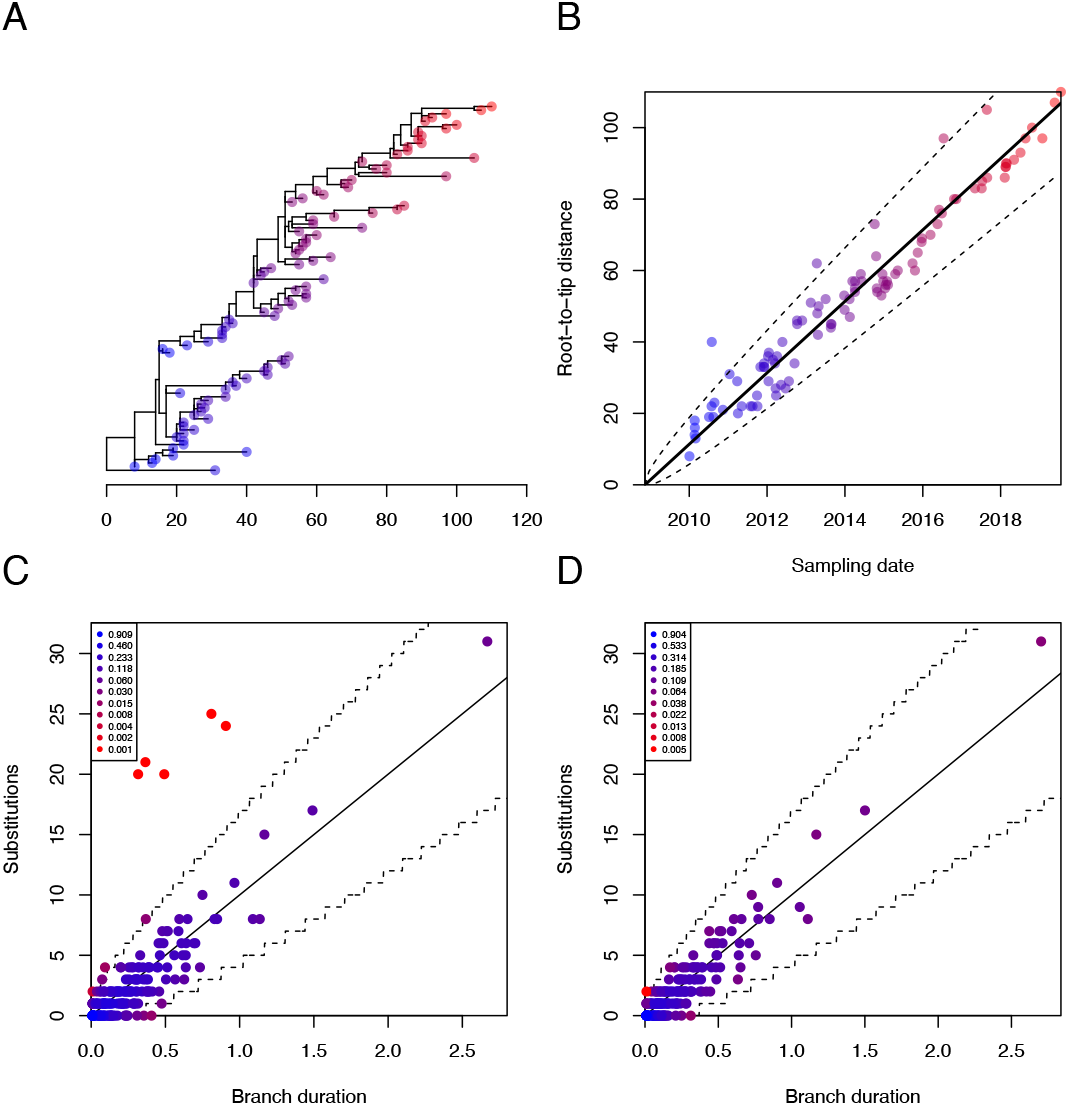
Example of diagnosis using outlier detection. (A) Simulated phylogeny with 5 outliers. (B) Root-to-tip regression analysis. (C) Distribution of substitutions per branch before removing outliers. (D) Distribution of substitutions per branch after removing outliers.

### Posterior predictive analysis

Posterior predictive assessment is a popular way to perform model diagnostics in Bayesian statistics Meng1994; Gelman et al.1996). It is frequently used for example in infectious disease epidemiology modeling (Didelot et al. 2017b; Whittles et al. 2017; Gibson et al. 2018). We therefore sought to apply it to the phylogenetic dating problem, using four carefully chosen summary statistics (see Methods): the mean of branch lengths, variance of branch lengths, maximum of branch lengths and the tree stemminess. The latter is defined as the sum of lengths of internal branches divided by the total sum of branch lengths, with high values usually indicating variations in population size (Fiala and Sokal 1985; Didelot et al. 2009).

A dated phylogeny was simulated including 100 leaves uniformly distributed between 2010 and 2020, under the heterochronous coalescent model (Drummond et al. 2002) with constant population size *N*_e_*g* = 1 year (Figure S1A). We applied the additive relaxed clock model ( Didelot et al. 2021) to this dated phylogeny, with mean clock rate *µ* = 10 substitutions per year and relaxation parameter *ω* = 5 (Equation 3). Consequently, some branches had many more or less substitutions compared to what would be expected under a strict clock model with *µ* = 10, and the probabilities of these branches under this model would be low (Figure S1B). Nevertheless, a root-to-tip regression seemed very satisfactory, with *R*^2^ = 0.94 and *p <* 10^−4^ for a date randomization test (Figure S2).

We applied BactDating (Didelot et al. 2018) to reconstruct the dated tree twice: first incorrectly using a strict clock model (Equation 1) and second correctly using the additive relaxed clock model (Equation 3). In the first case, the clock rate was estimated to be *µ* = 10.5 [9.4;11.6] and the root date 2008.6 [2008.1;2009.1]. In the second case, the clock rate estimated to be *µ* = 11.3 [8.8;14.1], the root date was 2008.9 [2007.7;2009.8] and the relaxation parameter was *ω* = 6.4 [4.2;8.9]. In both cases the values are approximately correct. BactDating includes a procedure to perform model comparison by computing the deviance information criterion (DIC) of each model ( Spiegelhalter et al. 2002). Here the strict clock model had a DIC of 1072.41, whereas the relaxed clock model had a DIC of 759.86, indicating strong support for the latter. This model comparison approach works well here since the data was generated here using the relaxed clock model. However, model comparison is not useful more generally to evaluate a single fit in absolute rather than relative terms. By contrast, the model diagnostic approach can identify poor model fits without the requirement of comparing to other models that may also be misspecified.

The posterior predictive check for the incorrect model diagnosed an issue on two summary statistics: the variance of the branch lengths and the tree stemminess (Figure 2A). On the other hand, when the correct model was used, the posterior predictive assessment did not detect any issue for any of the four summary statistics (Figure 2B). The posterior predictive assessment can also be applied more or less directly in the case where inference was performed using maximum likelihood methods ( Tsay 1992; Gelman et al. 1996, 2000; Lee et al. 2016), either by replacing the posterior samples with the point estimate or by building a pseudo-posterior (see Methods and next section).

**Figure 2:**
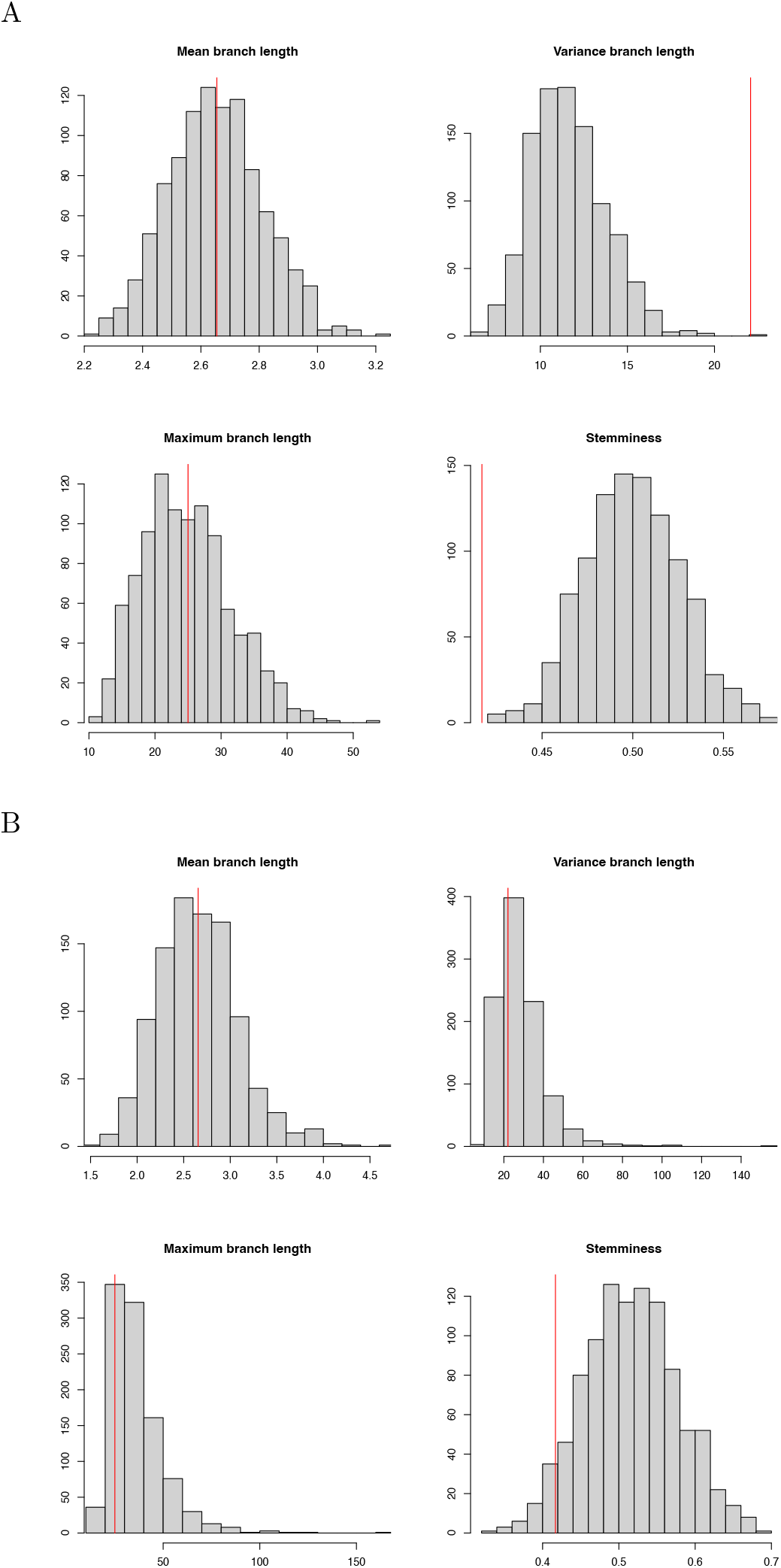
Example of posterior predictive analysis. (A) Inference using incorrect model. (B) Inference using correct model.

### Residual analysis

Another diagnostic approach is to consider the distribution of residuals after fitting a model. This methodology is especially reminiscent of regression models (Cox and Snell 1968; Dunn and Smyth 1996), but has also previously been applied more generally for example to epidemic models (Lau et al. 2014) or Hidden Markov Models (Zucchini and MacDonald 2009; Buckby et al. 2020). Here we adapt this approach to the problem of dating a phylogeny. Briefly, we consider for each branch the cumulative probability distribution of the number of substitutions given the branch duration. If the model is valid, these probabilities should be distributed as Uniform(0,1). However, it is diffcult to assess visually how close to zero or one a value needs to be in order to be an outlier. We therefore transform these probabilities into residuals with an expected distribution Normal(0,1), similar to the residuals used in regression models. This hypothesis can be evaluated using an Anderson-Darling simple hypothesis test (Lewis 1961). For more details see the Methods section.

Let us first consider the same two inferences as in the previous section, one from the incorrect strict clock model and one from the correct relaxed clock model. When looking at a single sample from the posterior using the incorrect model, the residuals for the branches were not distributed as expected (Figure 3A) and a QQ plot revealed significant deviation (Figure 3B). The Anderson-Darling test rejects the hypothesis of standard normality of the residuals (*p <* 10^−5^). By performing the same residual analysis on multiple samples from the posterior, we can construct a posterior distribution of p-values (Lau et al. 2014) in which all values were below 0.05, with a median below 10^−5^ (Figure S3A). The residuals for a single sample from the posterior using the correct model were approximately distributed as expected both when plotting them against their theoretical distribution (Figure 3C) and when constructing a QQ plot (Figure 3D). The Anderson-Darling test did not reject the hypothesis of standard normality of the residuals (*p* = 0.465). By repeating this residual analysis on multiple samples from the posterior we obtain a posterior distribution of p-values in which only 5.1% of them had p-values below 0.05, and the median was *p* = 0.513 (Figure S3B).

**Figure 3:**
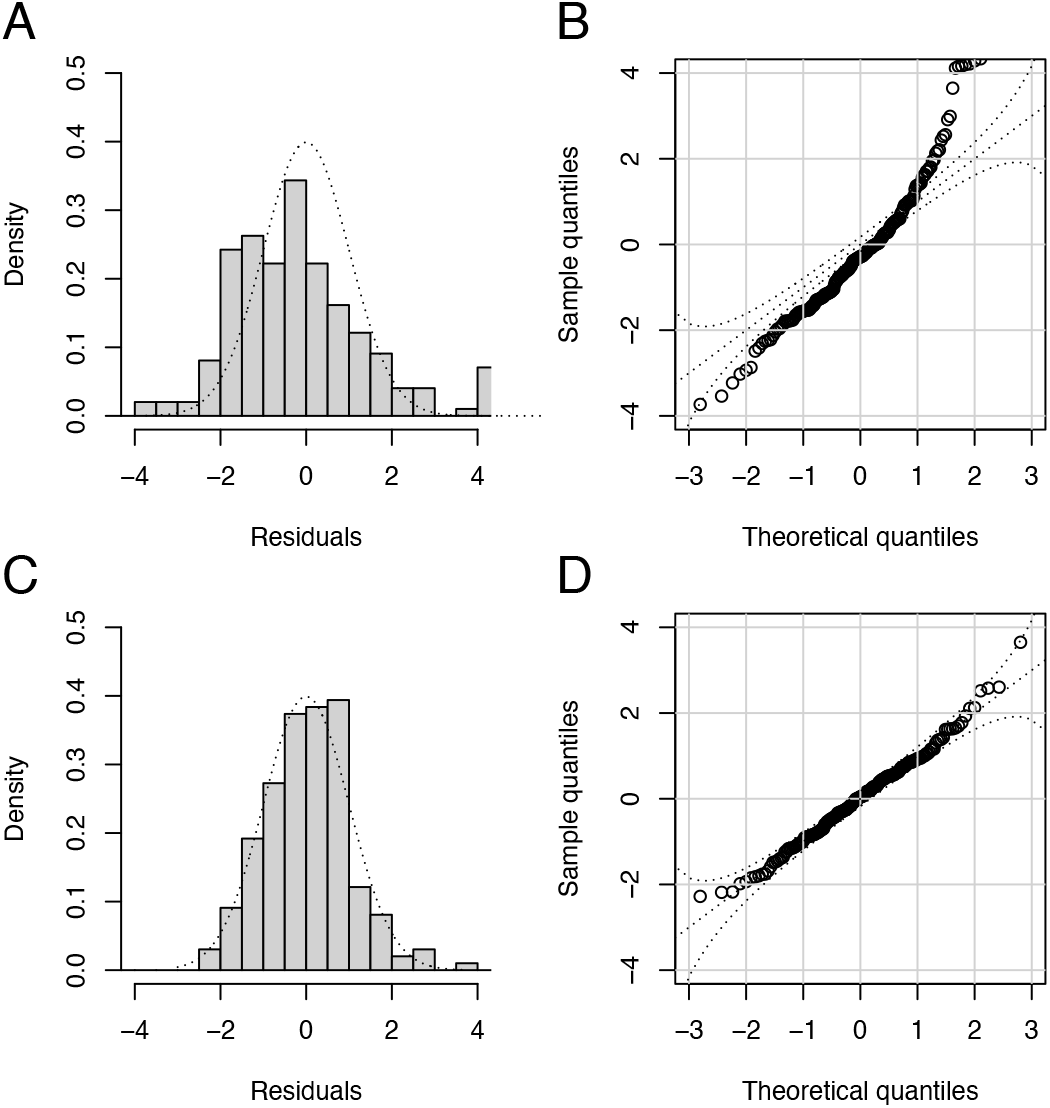
Example of residual analysis for a Bayesian inferred dated phylogeny. (A) Distribution of residuals after inference under a strict clock model. (B) QQ plot of residuals after inference under a strict clock model. (C) Distribution of residuals after inference under a relaxed clock model. (D) QQ plot of residuals after inference under a relaxed clock model.

We want to apply a similar residual analysis as before in the case of a point-estimated dated tree, for example using maximum-likelihood (ML) techniques. To illustrate this, we consider the dated tree in Figure 4A and simulate substitutions on each branch according to a strict clock model (Equation 1) with rate *µ* = 10 per year. The dated tree was inferred from this substitution data using treedater (Volz and Frost 2017) under the correct model, including estimation of the clock rate at *µ* = 9.32 and of the root date at 1972.83, which were close to the correct values. The likelihood of each branch is shown in Figure 4B. We computed the residuals for this ML tree as before and compared them the their expected Normal distribution, as shown in Figures 4C and 4D. It is visually clear that the residuals are underdispersed, and indeed the Anderson-Darling test had a p-value of 4 *·* 10^−4^. This underdispersion of residuals is expected when performing ML inference, and is a sign of overfitting a model that is overparametrised (Babyak 2004; Hawkins 2004). This can be illustrated using a simpler independently and identically distributed model for the branch lengths instead of a tree model (see Methods). This complicates the analysis of residuals compared the the previous Bayesian case, since we no longer have a straightforward expected distribution for the residuals. To remedy this problem, and bridge the gap between Bayesian and ML inference, we propose to generate an approximate Bayesian posterior sample centered around the ML inference (see Methods). The residuals for a single sample from this pseudo-posterior are shown in Figures 4E and 4F, from which it can be seen that they follow the expected Normal distribution. Indeed the Anderson-Darling test had a p-value of 0.42. As previously, we can compute such a p-value for all samples in the pseudo-posterior, which resulted in a posterior distribution of p-values (Figure S4) with 2.7% of values below 0.05, and the median p-value was *p* = 0.57.

**Figure 4:**
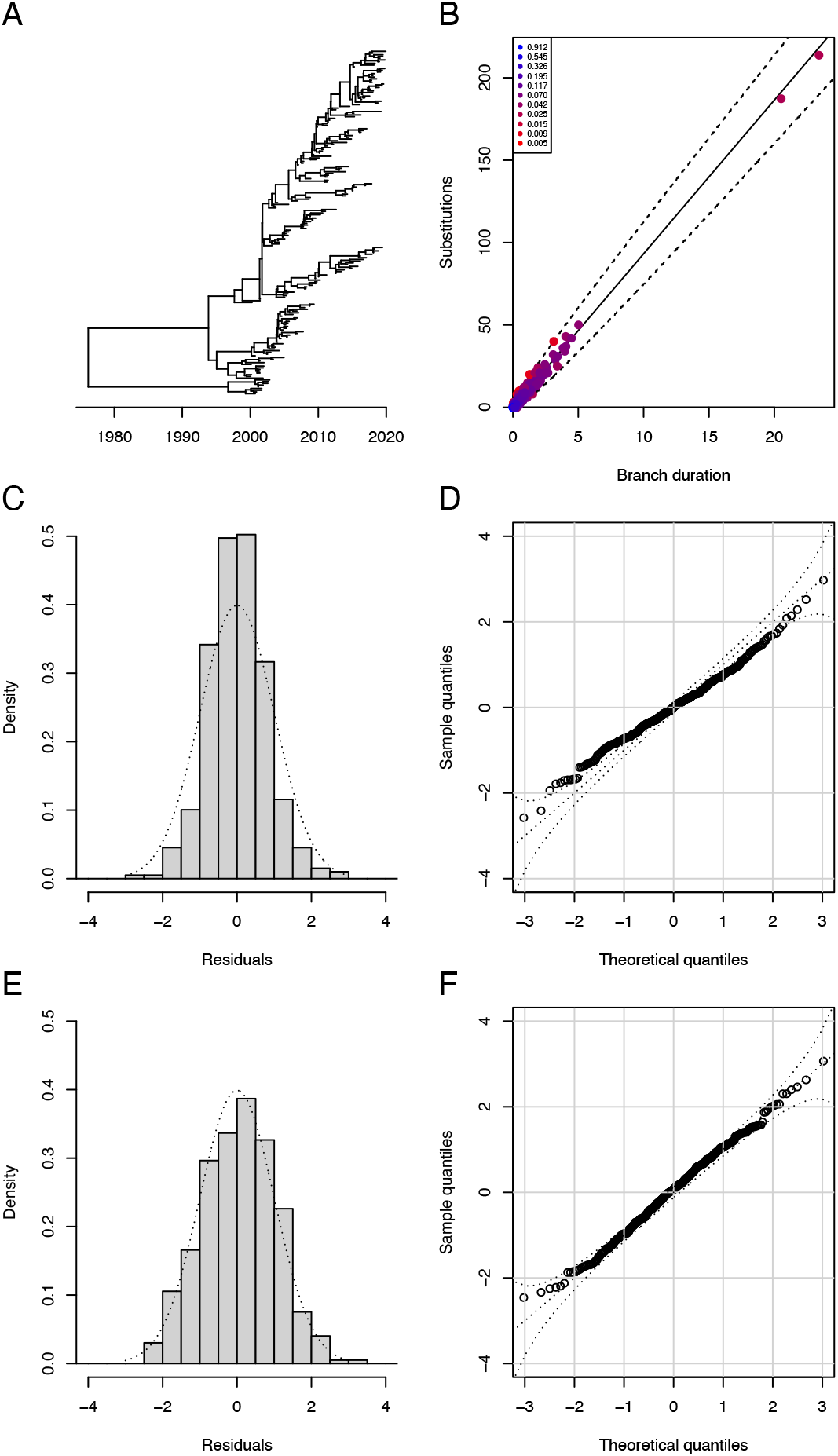
Example of residual analysis for a maximum-likelihood (ML) dated phylogeny. (A) Simulated dated phylogeny. (B) Likelihood of substitutions in the ML tree. (C) Distribution of residuals in the ML tree. (D) QQ plot of residuals in the ML tree. (E) Distribution of residuals in a tree from the pseudo-posterior. (F) QQ plot of residuals in a tree from the pseudo-posterior.

### Application to multiple simulated datasets

We simulated 100 datasets, each with 100 leaves uniformly distributed between 2010 and 2020, constant population size *N*_e_*g* = 1 year and a strict clock model with clock rate *µ* = 10 substitutions per year. For each dataset, we performed dating using five methods: BactDating (Didelot et al. 2018), treedater (Volz and Frost 2017), node.dating (Jones and Poon 2017), TreeTime (Sagulenko et al. 2018) and LSD (To et al. 2016). In each case the dating was performed under four conditions: with the root given, with the rate given, with both given and with neither given. Finally, for each inference we computed the p-value according to the posterior predictive analysis and residual analysis as described previously. We counted the number of times that the p-values were below 5% and the results are shown in Table 1. The number of problems diagnosed was low in all situations for BactDating, treedater and LSD, which is as expected since the same model was used for simulation and inference. On the other hand, both node.dating and TreeTime produced results for which issues were diagnosed using both the posterior predictive check and the residual analysis. However, these issues were detected only when the correct root was not provided, and irrespective of whether the rate was provided or not (Table 1). This suggests that these issues were often caused by a misidentification of the correct root in situations where the root was provided.

**Table 1:** Number of false positives found amongst a set of 100 replicates. Five different methods were used (BactDating, treedater, node.dating, TreeTime and LSD) and two different tests (posterior predictive check and residual analysis). Each method was applied in four different conditions: given the correct root, given the correct rate, given both or given neither.

| Method | Test | Given nothing | Given root | Given rate | Given root and rate |
| --- | --- | --- | --- | --- | --- |
| BactDating | PPcheck | 0 | 1 | 1 | 1 |
|  | Residuals | 0 | 1 | 0 | 1 |
| treedater | PPcheck | 0 | 1 | 0 | 1 |
|  | Residuals | 4 | 5 | 2 | 5 |
| node.dating | PPcheck | 23 | 3 | 28 | 2 |
|  | Residuals | 20 | 0 | 24 | 0 |
| TreeTime | PPcheck | 23 | 1 | 35 | 2 |
|  | Residuals | 35 | 3 | 40 | 2 |
| LSD | PPcheck | 0 | 1 | 0 | 1 |
|  | Residuals | 4 | 2 | 4 | 3 |

### Confounding effect of population structure

Population structure can have a confounding effect on dating analysis, especially when it is associated with systematic differences in sampling dates (Duchene et al. 2015; Murray et al. 2016; Tong et al. 2018). To explore whether this issue can be diagnosed, we simulated datasets with a population structure made of three components, each of which is sampled 50 times in three different years (cf Methods). These datasets are similar to the ones generated in a previous study (Murray et al. 2016) and we found evidence of confounding in about a third of datasets, but for illustration here we describe the results on a single example, for which the root of the tree existed in 1946 and a strict molecular clock (Equation 1) was used with rate 10 substitutions per year.

Figure 5A shows the simulated phylogeny. A root-to-tip regression analysis (Figure 5B) suggests a strong temporal signal, with no outliers. The correlation coeffcient was *R*^2^ = 0.95 and a date randomization test had a p-value below 10^−4^. However, the slope of this regression suggested a clock rate of 43 substitutions per year and the intersect with the x-axis suggested that the root existed in 2003. This is typical of what happens when population structure confounds the temporal signal, with an overestimation of the clock rate and an underestimation of the times to common ancestors (Murray et al. 2016). Note that this is the opposite of what happens when the temporal signal is weak, typically resulting in underestimation of clock rates and overestimation of times to common ancestors (Duchene et al. 2015), and therefore the tools previously proposed to ensure that the temporal signal is strong enough do not help diagnose this issue (Murray et al. 2016).

**Figure 5:**
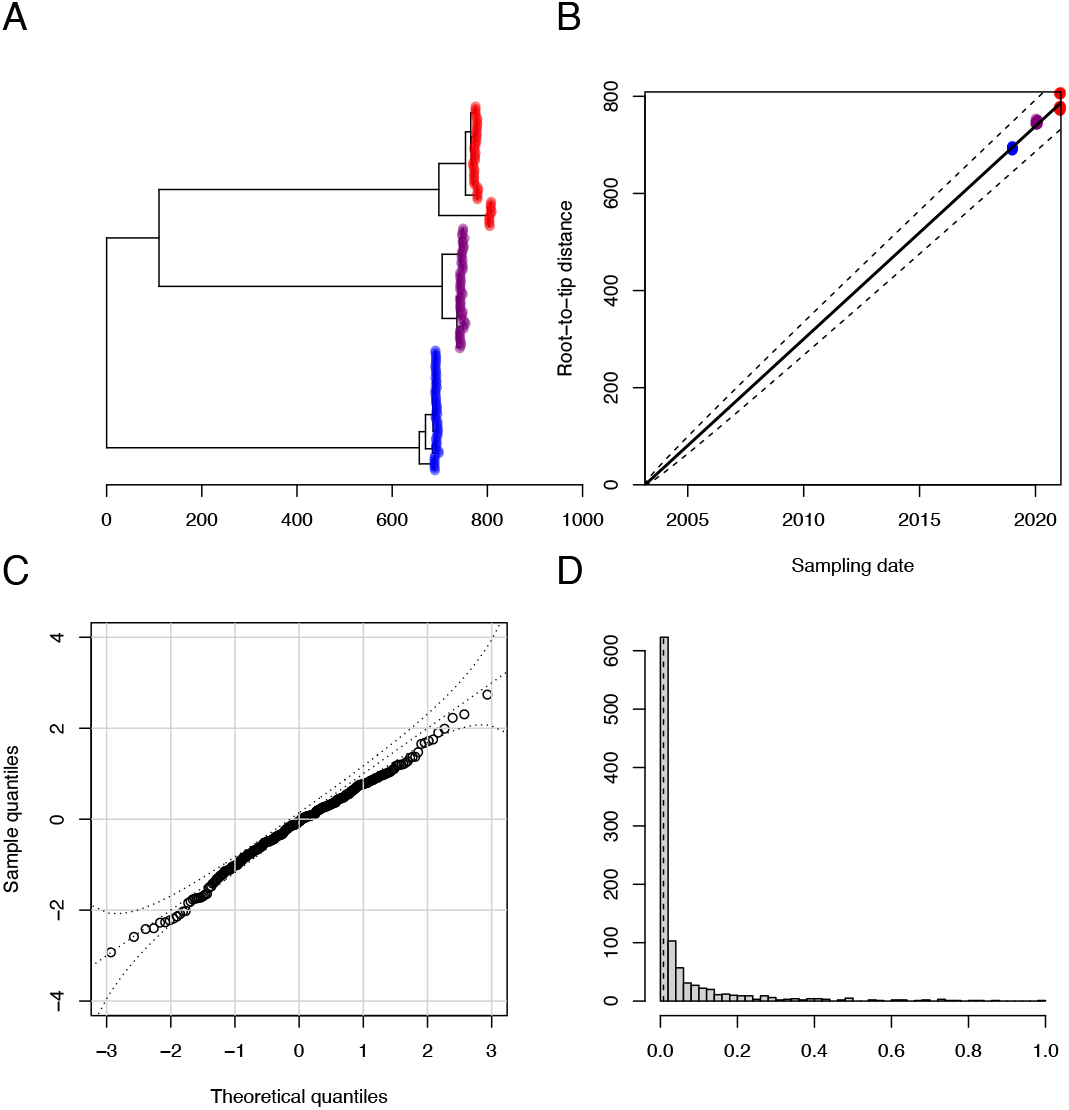
Example of diagnosis of the confounding effect of population structure. (A) Simulated phylogeny. (B) Root-to-tip regression analysis. (C) QQ plot of residuals for a single posterior sample. (D) Distribution of p-values from the residual analysis.

We ran BactDating (Didelot et al. 2018) on this dataset assuming the correct strict clock model, which estimated a clock rate of 16.4 [14.7-18.1] and a root date of 1976 [1971;1981]. Although these estimates are not as inaccurate as the ones quoted above for the root-to-tip analysis, the same issue is still clearly affecting the results with neither of the two credible intervals covering the ground truth values of 10 and 1946, respectively. The posterior predictive checks resulted in a p-value of 0.19 which is smaller than usually found for simulated datasets without structure, but not statistically significant. Figure 5C shows the QQ plot of residuals for a single posterior sample, for which the Anderson-Darling test rejects the hypothesis of standard normality of the residuals (*p* = 3 × 10^−3^). Repeating this test for all posterior samples resulted in the posterior distribution of p-values shown in Figure 5D, which has a median value of *p* = 4 × 10^−3^. The residual analysis was therefore successfully able to diagnose that there is an issue with the results.

### Application to Hepatitis B virus

We reanalysed a previously published dataset on Hepatitis B virus made of 137 whole genome sequences of aligned length 3,271bp (Patterson Ross et al. 2018). This included two ancient genomes from 1568 and 135 modern genomes sampled between 1963 and 2013. A phylogeny was built using PhyML (Guindon et al. 2010) and dating was performed using BactDating (Didelot et al. 2018) using the additive relaxed clock model (Didelot et al. 2021). The clock rate was estimated to be 9.02 × 10^−5^ per site per year (95% range between 7.82 × 10^−5^ and 1.04 × 10^−4^) and the root was estimated to have existed in 1162 (95% range between 1056 and 1247) as shown in Figure S5. However, applying diagnostics to this dating inference revealed that it should not be trusted. The posterior predictive checks had a p-value of 0.008 (Figure S6). The posterior distribution of residual p-values had a median of 2.04 × 10^−5^ (Figure S7). This result is consistent with a previous analysis of the same dataset which concluded that there was no temporal signal by comparing the likelihood when using the actual sampling dates versus constraining all dates to be equal (Duchene et al. 2020).

### Application to *Shigella sonnei*

We reanalysed a previously published dataset made of 155 Vietnamese whole genomes from the VN clade of *Shigella sonnei* (Holt et al. 2013). A previous analysis concluded that the additive relaxed clock model was well suited for analysing this dataset (Didelot et al. 2021). A root-to-tip regression analysis of this dataset looks especially satisfying, which could be taken as evidence of a good fit of a strict clock model (Figure S8). When fitting a strict clock model, the substitution rate was estimated to be 3.66 per genome per year (95% range between 3.29 and 3.98) and the date of the most recent common ancestor was 1982 (95% range between 1979 and 1985). The posterior predictive checks did not reject this inference (p-value 0.37), but the analysis of residuals rejected it, with a median p-value of 0.0019. Indeed in a posterior sampled dated tree, (Figure 6A) there are 12 branches with residual probability below 0.01 (Figure 6B). We repeated the analysis using the additive clock model, which gave similar estimates for the clock rate (3.75 with range 3.33 to 4.16) and the root date (1982 with range 1979 to 1985). This time however neither the posterior predictive analysis (Figure S9) nor the residual analysis (Figure S10) rejected the result, with p-values of 0.49 and 0.26, respectively. A posterior sampled dated tree (Figure 6C) looks very similar to the one estimated using the strict clock model (Figure 6A) but there is now only a single branch with residual probability below 0.01 (Figure 6D), which is not unexpected in a tree containing 308 branches. This result therefore agrees with the previous analysis that had found the additive relaxed clock model to be best based on model comparison (Didelot et al. 2021), and goes a step further by validating this model in absolute rather than relative terms.

**Figure 6:**
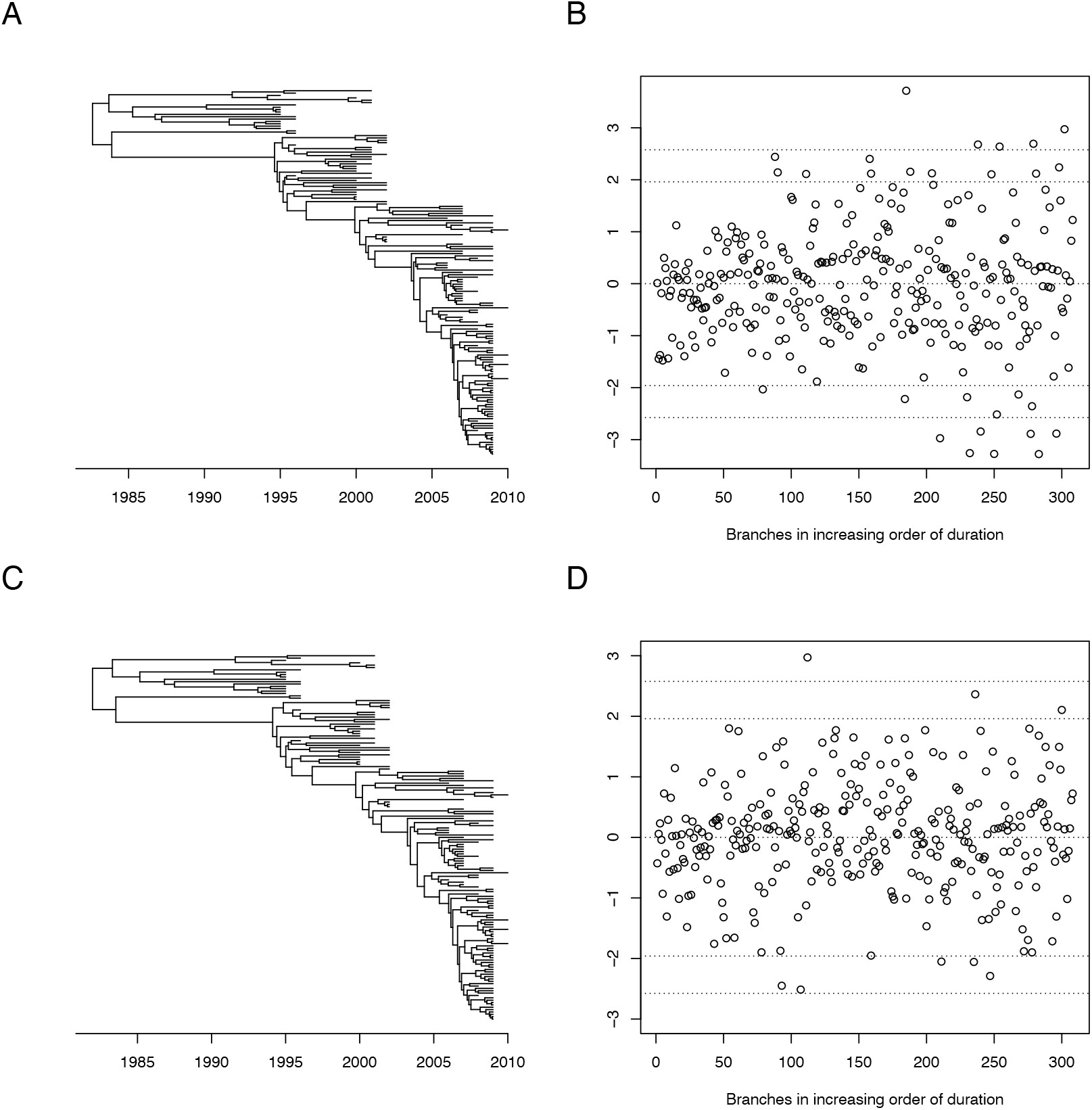
Analysis of the *Shigella sonnei* dataset. (A) Dated tree estimated when using a strict clock model. (B) Residual distribution when using a strict clock model. (C) Dated tree estimated when using an additive relaxed clock model. (D) Residual distribution when using an additive relaxed clock model.

### Application to *Streptococcus pneumoniae*

We reanalysed a previously published dataset made of 238 whole genomes from the PMEN1 lineage of *Streptococcus pneumoniae* sampled between 1984 and 2008 (Croucher et al. 2011). The original study found that recombination occurred frequently in this dataset, to the point that it made the temporal signal unclear (Croucher et al. 2011). Another analysis of the same dataset found that correcting for recombination using Gubbins (Croucher et al. 2015) resulted in more accurate and precise estimates of the ancestral dates (Didelot et al. 2018). We analysed this dataset again first without correction for recombination (Figure 7A). The branches in the resulting dated trees show only a weak correlation between duration and number of substitutions, which can only be explained by a highly relaxed molecular clock (Figure 7B). The analysis of residuals rejected this result, with a median of the posterior distribution of p-values of 0.006. On the other hand, the analysis after correction for recombination (Figure 7C) results in a tree with a much higher correlation between branch durations and numbers of substitutions (Figure 7D) so that the relaxed clock model has a relaxation parameter 5 times lower than in the analysis without correction for recombination (1.04 vs 5.68). The analysis of residuals did not diagnose an issue with this result (Figure S11), with a median of the posterior distribution of p-values of 0.25. This result underlines the importance of accounting for recombination when dating recombinant bacterial lineages (Didelot and Parkhill 2022).

**Figure 7:**
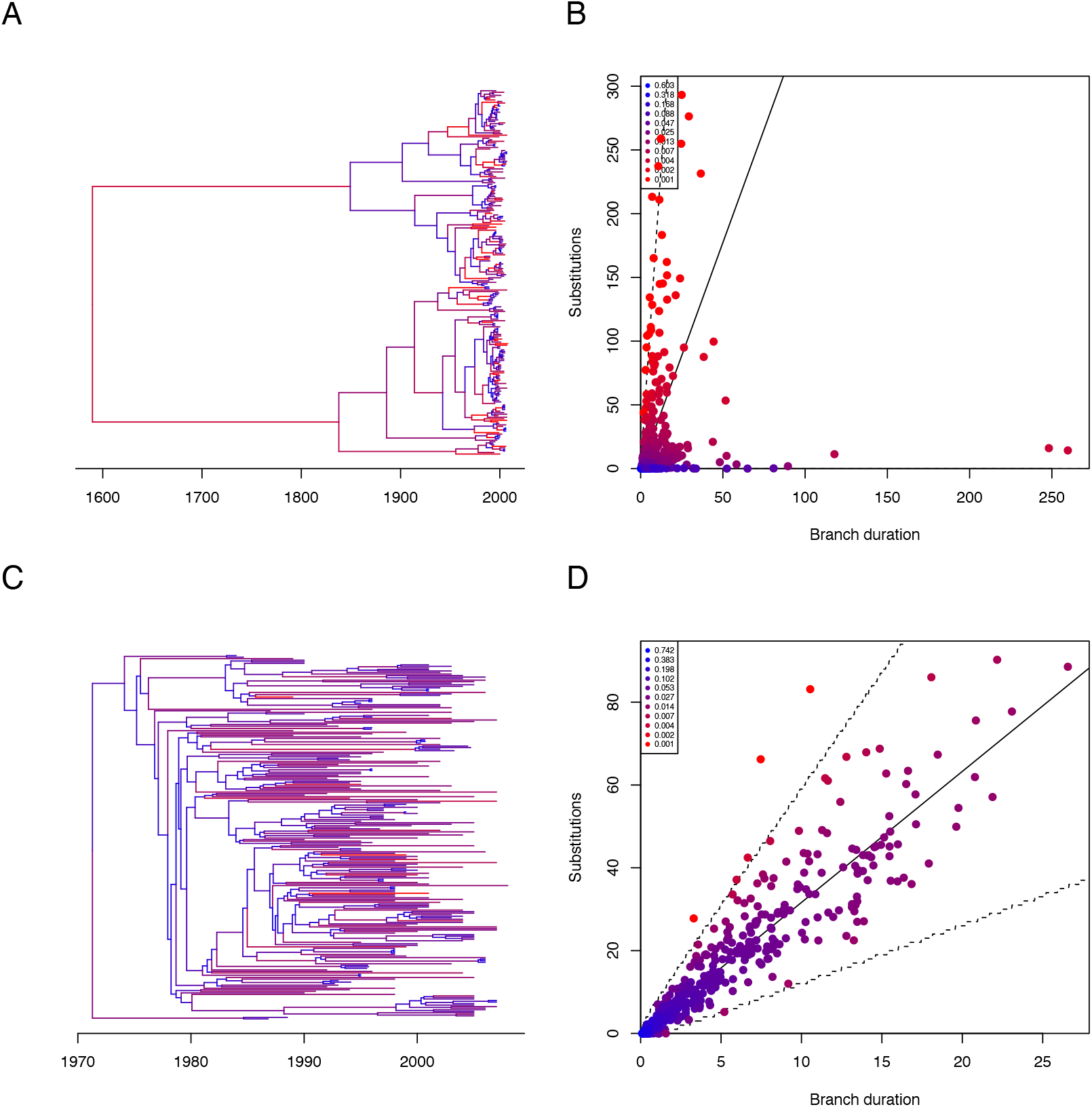
Analysis of the *Streptococcus pneumoniae* dataset. (A) Dated tree estimated without correction for recombination. (B) Likelihood of branches in the analysis without correction for recombination. (C) Dated tree estimated with correction for recombination. (D) Likelihood of branches in the analysis with correction for recombination.

## DISCUSSION

In this study we have investigated the use of model diagnostic methods after inference of a dated phylogeny in microbial population genetics. We have briefly reviewed the use of outlier detection analysis which has been proposed previously (Rambaut et al. 2016). We have also adapted to this setting two methods that are frequently used in other statistical areas, namely posterior predictive checking (Meng 1994; Gelman et al. 1996) and residual analysis (Dunn and Smyth 1996; Lau et al. 2014). The posterior predictive checking approach requires to select summary statistics informative on whether the model fit is correct or not. We use four summary statistics selected to represent different aspects of the phylogeny, and we found them to be a useful choice in practice but we do not claim that they are in any way optimal. There are other options that could be explored such as the coeffcient of variation of rates in the tree (Volz and Frost 2017) or the phylogenetic tree shape (Colijn and Plazzotta 2018). On the other hand, using more summary statistics could reduce the statistical power of the test due to the need to correct for multiple testing.

We found the residual analysis approach to be highly promising in terms of specificity and sensitivity when applied to simulated datasets. When applied to real datasets, we found that this method can help diagnose a wide range of issues, such as lack of temporal signal, inappropriate molecular clock model or distorting effect of recombination. Standard phylogenetic methods do not account for recombination, which can distort branch lengths in particular (Hedge and Wilson 2014). Phylogenetic methods have been developed specifically to resolve this issue, such as Gubbins (Croucher et al. 2015) or ClonalFrame (Didelot and Wilson 2015), but are known to be imperfect and detect and correct only the more impactful recombination events. Model diagnostics are therefore useful in ensuring that inferred dated phylogenies can be trusted, and should be considered an essential part of the pipeline for the large-scale analysis of bacterial genomes (Didelot and Parkhill 2022).

We have focused on diagnostics for dated phylogenies constructed using methods that date the nodes of an undated phylogeny, for example BactDating (Didelot et al. 2018), TreeTime (Sagulenko et al. 2018) or treedater (Volz and Frost 2017). However, similar diagnostics could also be applied to methods that build a dated phylogeny directly from the alignment, such as BEAST (Suchard et al. 2018). The posterior predictive approach (Meng 1994; Gelman et al. 1996) would need to be modified, so that the input alignment is compared with simulated alignments based on the posterior distribution of dated phylogenies and molecular clock model parameters. This is straightforward in principle, but in practice it would require the generation of many large simulated datasets, and comparisons between summary statistics of the alignments rather than the phylogenies as we did here. Finding informative summary statistics for alignments is likely to be more diffcult than for phylogenies. The residual analysis approach we proposed would also need adapting when dating does not start with an undated phylogeny. A separate undated phylogeny could be constructed from the same alignment, but this may not have the same topology as the inferred dated phylogeny so that residuals are not as easy to compute. Alternatively, an ancestral state reconstruction given a dated tree could be performed (Pupko et al. 2000; Joy et al. 2016) so that substitutions can be counted on the branches and residuals calculated accordingly.

In conclusion, we have found that it is possible to diagnose a wide range of potential issues that can arise when dating a phylogeny. When a dating inference is found to be invalid, it means that at least one of the assumptions made by the underlying model is incorrect. Conversely, the fact that diagnostic tests are passed means that the underlying assumptions cannot be rejected but does not guarantee that they are correct either (since “absence of evidence is not evidence of absence”). As in other branches of statistical inference, model diagnostics provides insights that are complementary with the use of model comparison. We recommend that model diagnostics and model comparison should be used consistently in the growing number of microbial population genetic studies that rely on the reconstruction of dated phylogenies.

## MATERIALS AND METHODS

### Molecular clock models

The molecular clock model determines the distribution of number of substitutions *l*_*i*_ on a branch of the dated tree with duration *d*_*i*_. We consider four types of molecular clock models, for each combination of whether the values *l*_*i*_ are discrete or continuous and whether the clock model is strict or relaxed. In the discrete strict clock model (Zuckerkandl and Pauling 1962) with rate *µ*, substitutions occur on the branches as a Poisson process with rate *µ* and therefore:

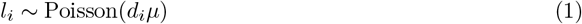

A continuous version of the strict clock model can be formed based on a Gamma process with the same mean and variance (Didelot et al. 2021):

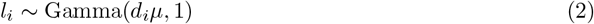

Strict clock models are based on the assumptions that the substitution rate is constant throughout the branches of the tree, but this is not always true, in which case a relaxed clock model can be used that allows the rate to vary (Drummond et al. 2006). In particular here we use the additive relaxed clock model (Didelot et al. 2021), in which *µ* is the mean clock rate and *ω* determines how much this rate varies on the branches. The discrete version of this model is given by:

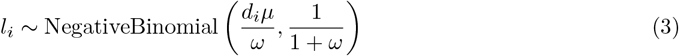

A continuous additive relaxed clock model can again be defined by considering a Gamma process with the same mean and variance as the discrete version:

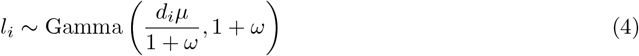

Note that throughout this article Gamma distributions are parametrized by shape and scale and Negative Binomials by number of successes and probability of success. In the four models we have that the mean of *l*_*i*_ is equal to *d*_*i*_*µ*. The variance of *l*_*i*_ is equal to its mean in the two strict clock models, and equal to its mean times (1 + *ω*) in the two relaxed clock models.

### Posterior predictive checks

When applied to the phylogenetic dating problem, posterior predictive checking requires the simulation of many undated phylogenies from the posterior sample of dated phylogeny, and comparing them to the observed undated phylogeny from which inference was performed. Simulation is done using the same clock model as was used for inference (Equations 1 to 4) and using the posterior inferred parameters. Posterior predictive assessment can also be performed following inference from a maximization method, using the single inferred dated tree and parameters as starting point for all simulations. The comparison of simulated and observed phylogenies is done on the basis of summary statistics, and here we used the following four: mean of branch lengths, variance of branch lengths, maximum of branch lengths and stemminess (Fiala and Sokal 1985). For each summary statistic, an empirical p-value is computed representing how extreme the observed phylogeny is compared to the set of simulated ones. These p-values can then be combined into a single p-value while controlling for multiple testing. We used a false discovery rate (FDR) correction (Benjamini and Hochberg1995) although other options would also be possible for example using a harmonic mean p-value (Wilson 2019).

### Computation and analysis of residuals

We want to diagnose a dated phylogeny 𝒟 by comparison with an undated phylogeny ℒ. We start by considering the case where the dated phylogeny 𝒟 is a single sample from the posterior distribution *p*(.|ℒ) obtained for example using BactDating (Didelot et al. 2018). Let *d*_*i*_ be the duration of a given branch in *D* and *l*_*i*_ be the number of substitutions on the corresponding branch of ℒ, that is the branch that separates the leaves in the same way. There is a unique corresponding branch in ℒ for all branches in 𝒟 except for the two branches *a* and *b* connected to the root of 𝒟 for which there is only a single corresponding branch *x*. We therefore split the substitutions on *x* proportionally between the two branches *a* and *b* by defining:

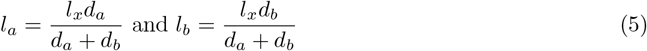

The distribution of *l*_*i*_ given *d*_*i*_ is given by the molecular clock model. Let us for now consider that the distribution is continuous (as in Equations 2 and 4) and we will return later to the discrete case (as in Equations 1 and 3). Instead of a specific model, we consider the general case where *F*_*i*_(*l*_*i*_) is the cumulative distribution function of *l*_*i*_ given *d*_*i*_. Let *u*_*i*_ denote the uniform residual for the observation *l*_*i*_, defined as:

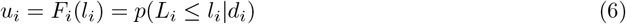

If the inference is valid, then the uniform residual *u*_*i*_ should be distributed as Uniform(0,1), because for any random variable *X* with cumulative distribution function *F* we have that *U* = *F* (*X*) is Uniform(0,1). We can then define the normal residuals *n*_*i*_, analogous to the residuals commonly used in regression models (Cox and Snell 1968; Dunn and Smyth 1996). The normal residuals are obtained by transforming the uniform residuals with the inverse of the cumulative distribution function Ф of a Normal(0,1) random variable:

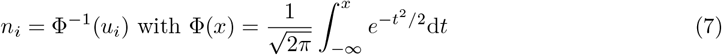

If the inference is valid, then the normal residuals *n*_*i*_ should be distributed as Normal(0,1) which is more convenient to work with than the Uniform(0,1) for uniform residuals. The uniform and normal residuals above can be computed directly when the clock model is continuous (Equations 2 and 4) but when the clock model is discrete (Equations 1 and 3) we need to make the following adjustment (Dunn and Smyth 1996; Brockwell 2007; Lau et al. 2014):

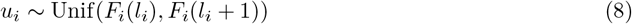

After computation of the uniform residuals *u*_*i*_ and normal residuals *n*_*i*_ for each branch, we use several methods to assess the validity of the dated phylogeny inference. The uniform residuals *u*_*i*_ can be plotted as a histogram to compare their distribution with the theoretical Uniform(0,1) distribution, but as noted above this can be diffcult to interpret. We therefore prefer to use the normal residuals *n*_*i*_ which can be plotted as a histogram to compare their distribution with the theoretical Normal(0,1). A quantile-quantile plot (QQ plot) can be used to compare the distribution of the residuals to their theoretical distribution. A p-value can be computed to assess that the normal residuals are distributed as expected. The most commonly used and powerful test of normality is the Shapiro-Wilk test (Razali and Wah 2011), but it is a composite test for when the mean and variance are unknown, whereas here we know that the normal residuals follow the standard normal distribution with mean 0 and variance 1. We therefore use the Anderson-Darling simple hypothesis test (Lewis 1961) as implemented in the DescTools R package based on previously published code (Marsaglia and Marsaglia 2004). Application of the Anderson-Darling test on the normal residuals *n*_*i*_ defined in Equation 7 against a Normal(0,1) distribution is exactly equivalent to application of the Anderson-Darling test on the uniform residuals *u*_*i*_ from Equation 7 against a Uniform(0,1) distribution.

We have described the diagnostics procedure above as if there was a single posterior sample 𝒟 to diagnose, whereas there would typically be multiple samples from this posterior available for example from running a Markov Chain Monte-Carlo method (Didelot et al. 2018). However, the same computation and analysis of residuals can be performed for each sample as described above. Each statistical test will return a separate p-value and these can be combined to form a posterior distribution of p-values (Streftaris and Gibson 2012; Lau et al. 2014; Gibson et al. 2018). From this posterior distribution of p-values we can compute various summaries to measure the validity of the inference, for example the proportion of p-values that are below 0.05 (Lau et al. 2014). We also often report the median of the posterior sample of p-values, since it can be interpreted more directly as a p-value.

### Pseudo-posterior sampling given a point estimate

We now consider the case where the dated phylogeny 𝒟 that we wish to diagnose was not sampled from the posterior. Instead it may be a point estimate, for example the result of maximum likelihood estimation (Volz and Frost 2017; Sagulenko et al. 2018) or a summary tree built from a posterior sample (Heled and Bouckaert 2013) but for which the posterior sample itself is not available. In this case we propose to first generate approximate samples from the posterior before residuals can be computed as described above. If the residuals were computed directly from the point estimate, they would not follow the same distribution as in Equations 6 and 8. To explain and illustrate this, let us first consider the discrete strict clock model (Equation 1) with known mutation rate *µ* = 1 and that the true branch durations *d*_*i*_ are independent and identically distribution as:

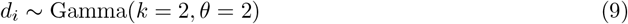

Let 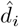 be a branch length in 𝒟, on which there are *l*_*i*_ substitutions in the undated phylogeny ℒ. The true residuals of each branch are distributed as expected (Figure S12A) and so are the residuals based on a posterior sample of *d*_*i*_ given *l*_*i*_ (Figure S12B). If 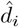 is a maximum likelihood estimate of *d*_*i*_ then 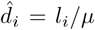. The residuals of the data points *l*_*i*_ against the estimates 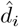 are underdispersed, because the maximum likelihood estimates are overfitted (Figure S12C). It is however possible in this case to recover the exact correct residuals by sampling from the posterior. By conjugacy of the Gamma prior and Poisson likelihood, we can deduce that the posterior of *d*_*i*_ is:

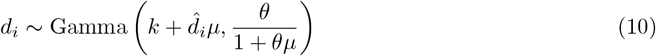

We can simulate from this distribution to get a posterior sample *d*_*i*_, from which we can then compute the residuals as described previously which will be distributed as in Equations 6 and 8 (Figure S12D).

The case described above works exactly but is idealised for two reasons. Firstly the branch durations *d*_*i*_ are neither independent nor identically distributed, but rather related through each other via the coalescent process. Secondly the clock is not usually strict with a known rate so that a posterior would not be analytically available. However, we can follow a similar idea of generating a pseudo-posterior sample centered on the given point estimate 𝒟. To do so, we perform a short run of BactDating (Didelot et al. 2018) for the input phylogeny ℛ with branch lengths equal to the lengths in 𝒟 multiplied by the clock rate 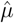 estimated when the dated tree was inferred. If this estimate is not available, a simple maximum likelihood estimator can be used instead:

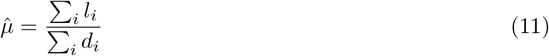

Since the branch lengths in ℛ are continuous, inference is performed under the continuous version of the strict clock model (Equation 2). The clock rate is fixed equal to its previous estimate 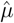, and the coalescent rate α is fixed equal to either its previous estimate (if available) or the maximum likelihood estimator:

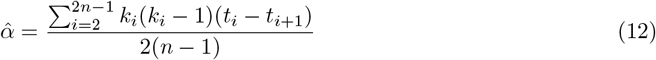

Note that this corresponds to the mean of the posterior distribution of α assuming an improper InvGamma(0, ∞) prior on α so that posterior is:

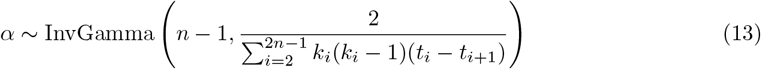

The posterior sample returned by BactDating can be thought of as a pseudo-posterior centered on the point estimate 𝒟, from which residuals can be computed as in the Bayesian case. An alternative would be to use a parametric bootstrap (Efron 2012). Generating a thousand bootstrapped datasets is easy simply by application of one of the molecular clock models (Equations 1 to 4) to the point estimate 𝒟, but inference would then have to be performed for each bootstrapped dataset which is not computationally attractive for the size of datasets considered here.

### Data simulation

The unstructured datasets were simulated by first sampling from the heterochronous coalescent model (Drummond et al. 2002) and then applying one of the molecular clock models (Equations 1 to 4). We tested several approaches to the simulation of confounding structured datasets (Murray et al. 2016), including using DetectImports (Didelot et al. 2023b) and using Master (Vaughan and Drummond 2013) to simulate under the structured coalescent model (Nordborg 1997), but these methods would typically not cause enough confounding effect for our purposes. We therefore implemented our own simulation method using mlesky (Didelot et al. 2023a) to simulate each population component genealogy 𝒢 under a coalescent model with a non-constant population size *N* (*t*), so that its probability follows:

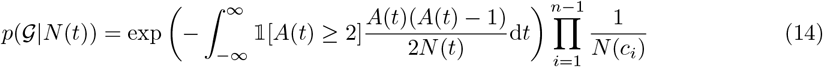

where *A*(*t*) represents the number of lineages at time *t* and *c*_*i*_ represents the times of the nodes. The size of the *j*-th population component followed a previously studied model of clonal expansion (Helekal et al. 2021):

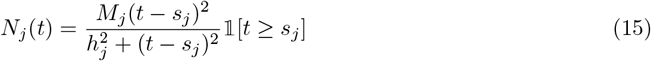

Each population component starts at time *s*_*j*_ with size *N* (*s*_*j*_) = 0 and grows logistically up to its maximum *N*_*j*_(∞) = *M*_*j*_, with *h*_*j*_ being the time taken to reach half of this since *N*_*j*_(*s*_*j*_ + *h*_*j*_) = *M*_*j*_*/*2.

### Implementation

We implemented the analytical methods described in this paper in a new R package entitled *DiagnoDating* which is available at https://github.com/xavierdidelot/DiagnoDating for R version 3.5 or later. *DiagnoDating* provides a joint interface to run analyses using LSD (To et al. 2016), node.dating (Jones and Poon 2017), treedater (Volz and Frost 2017), BactDating (Didelot et al. 2018) or TreeTime (Sagulenko et al. 2018), and to compute all the diagnostic methods we described.

All code and data needed to replicate the results are included in the “reproducibility” directory.

