## Supplementary Material for "Diagnostics of dated phylogenies in microbial population genetics"

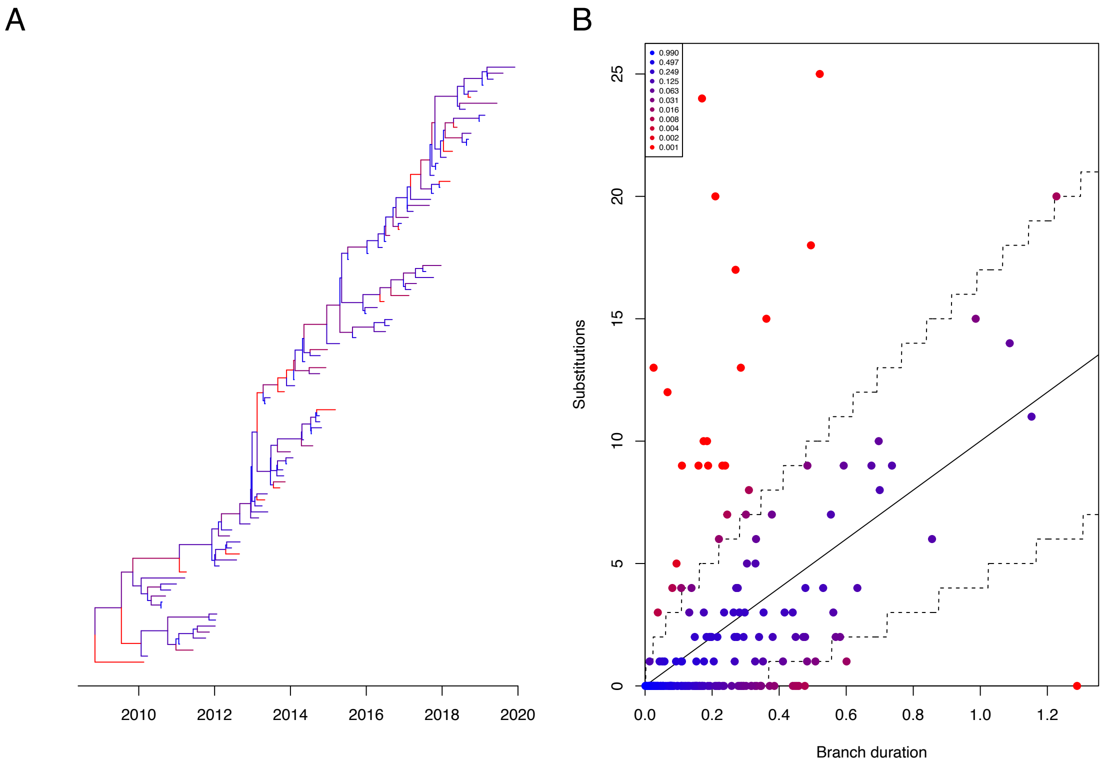


Figure S1: (A) Dated tree used as motivating example. (B) Likelihood of branches under a strict clock model with rate 10.

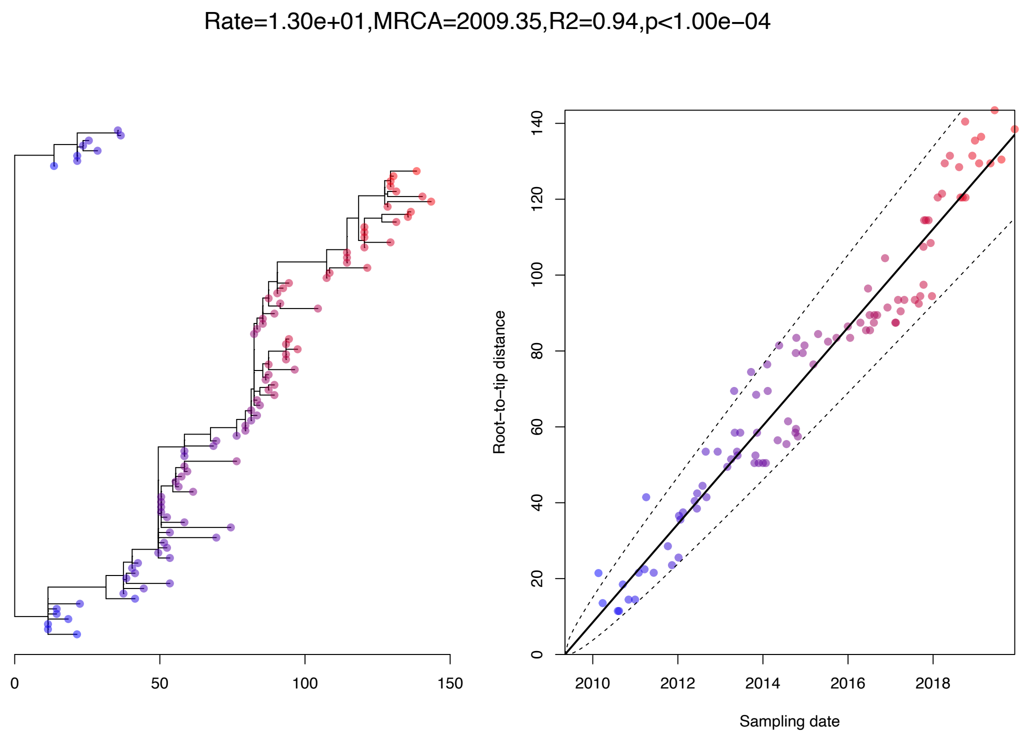


Figure S2: Root-to-tip regression analysis for the motivating example.

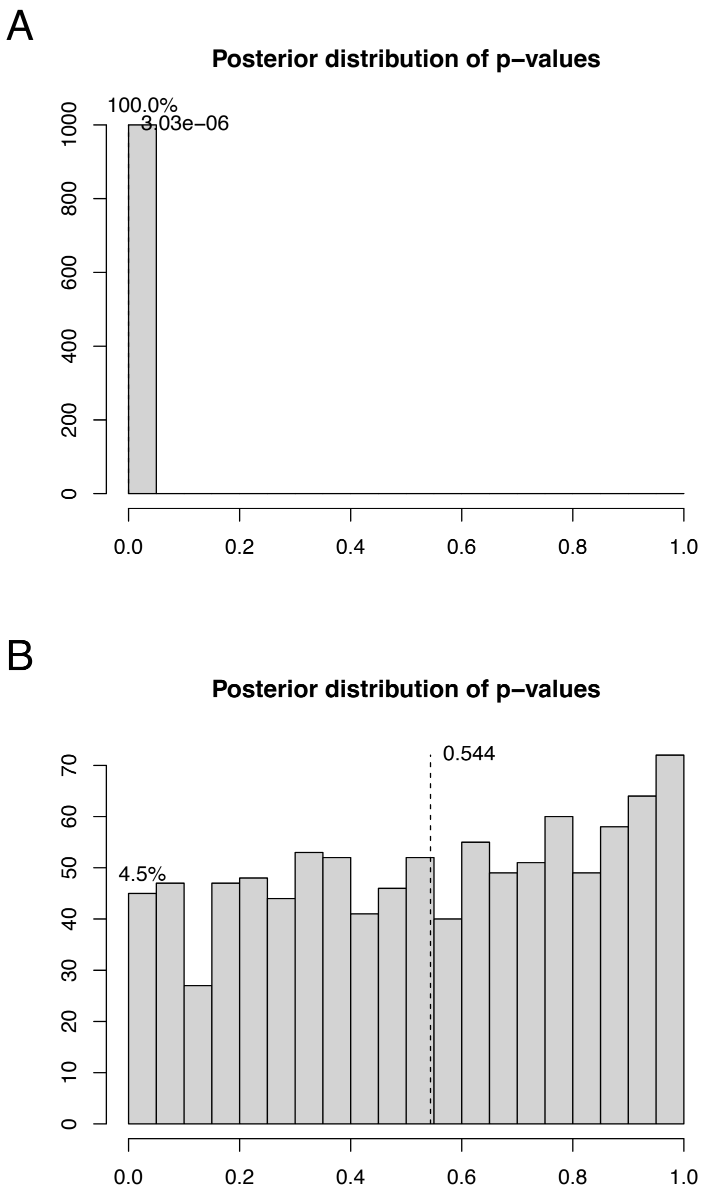


Figure S3: Posterior distribution of p-values for inference under the strict clock model (A) and ARC model (B).

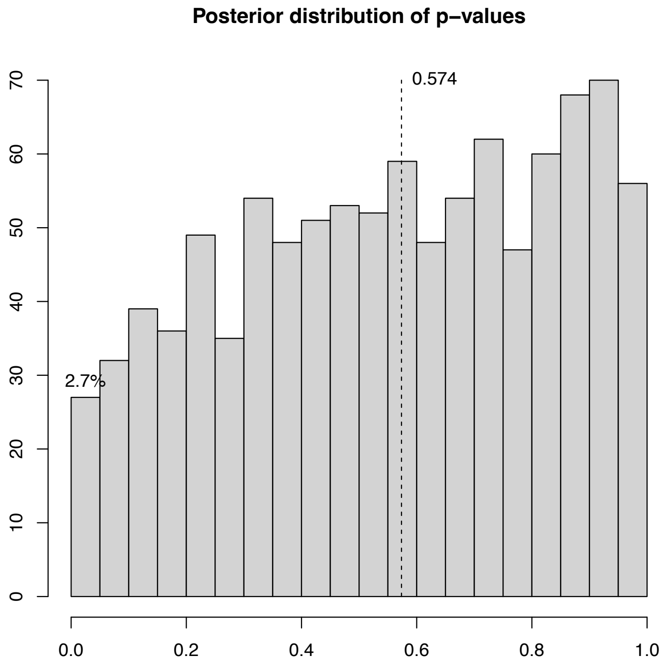


Figure S4: Posterior distribution of p-values for the pseudo-posterior based on ML inference.

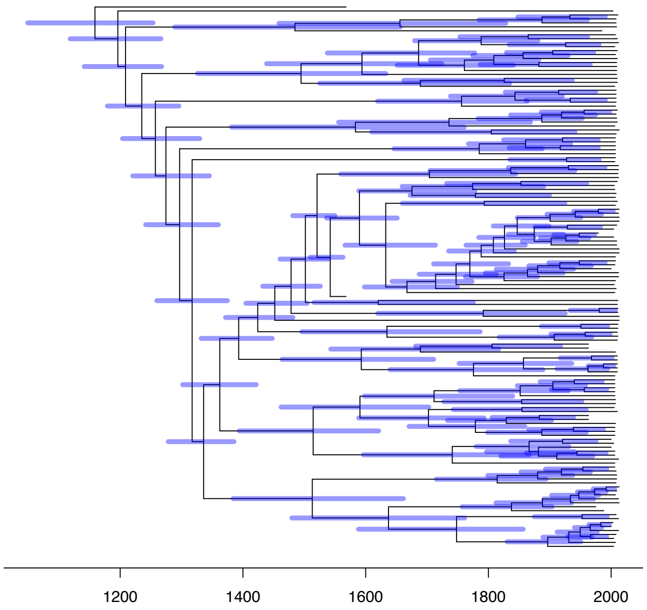


Figure S5: Dated phylogeny estimated for the HBV dataset.

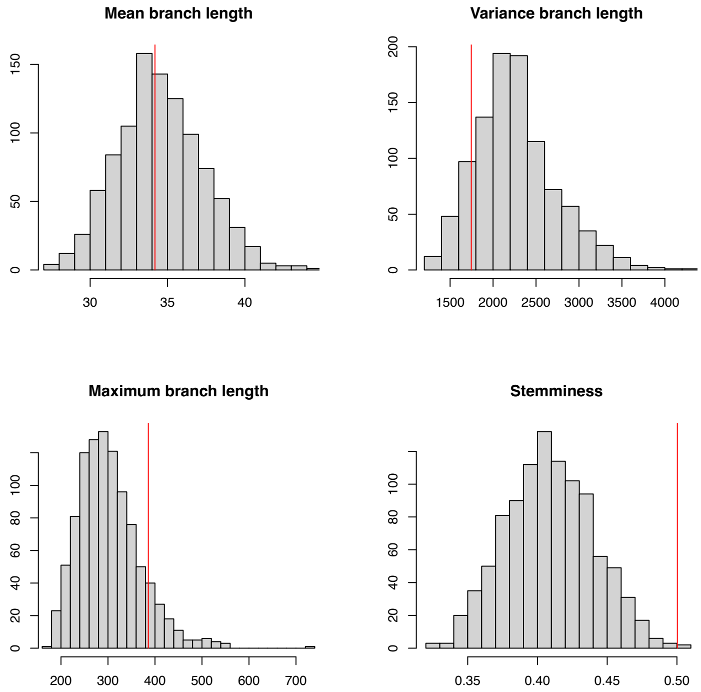


Figure S6: Posterior predictive checks for the HBV application.

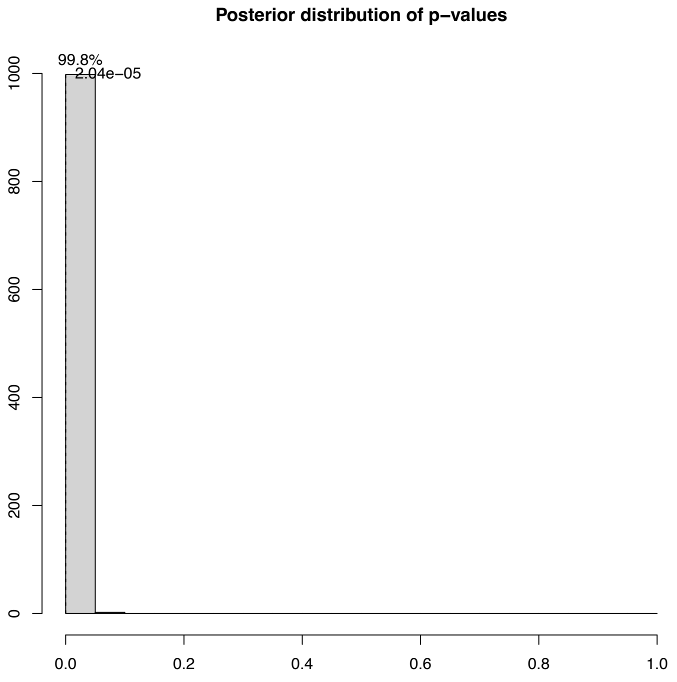


Figure S7: Posterior distribution of residual p-values for the HBV application.

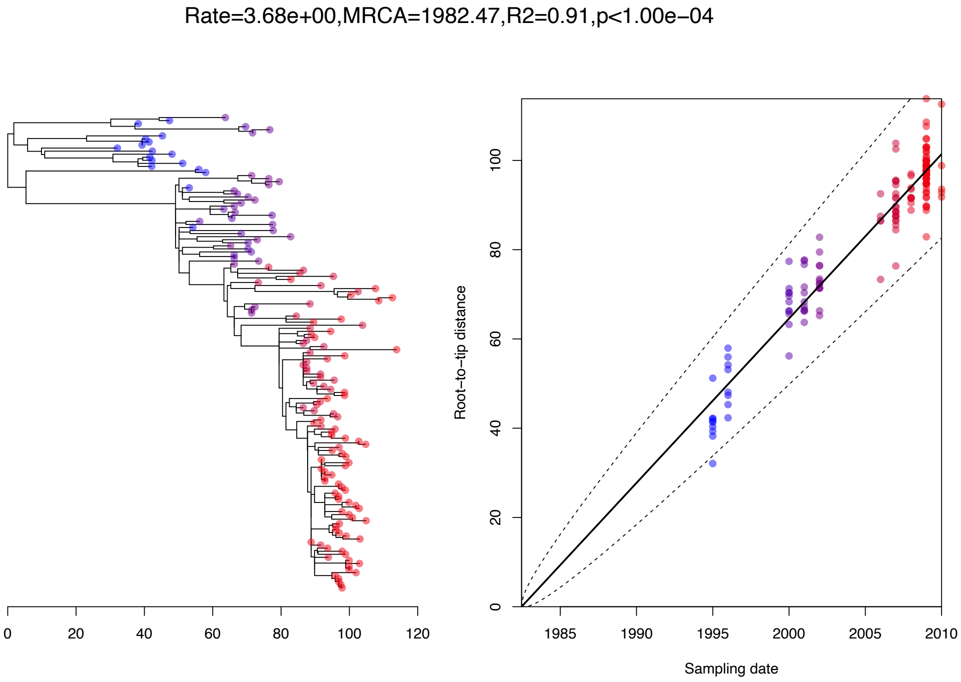


Figure S8: Root-to-tip analysis for the Shigella dataset.

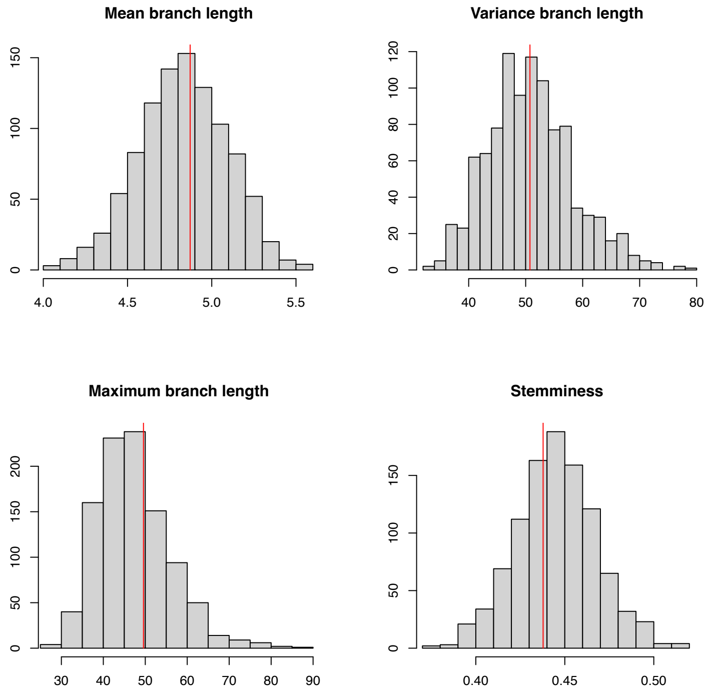


Figure S9: Posterior predictive checks for the Shigella application.

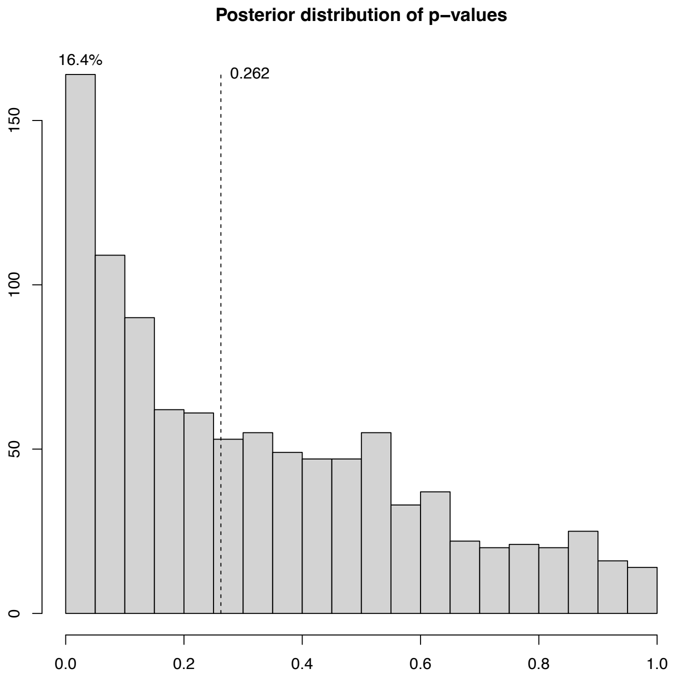


Figure S10: Posterior distribution of residual p-values for the Shigella application.

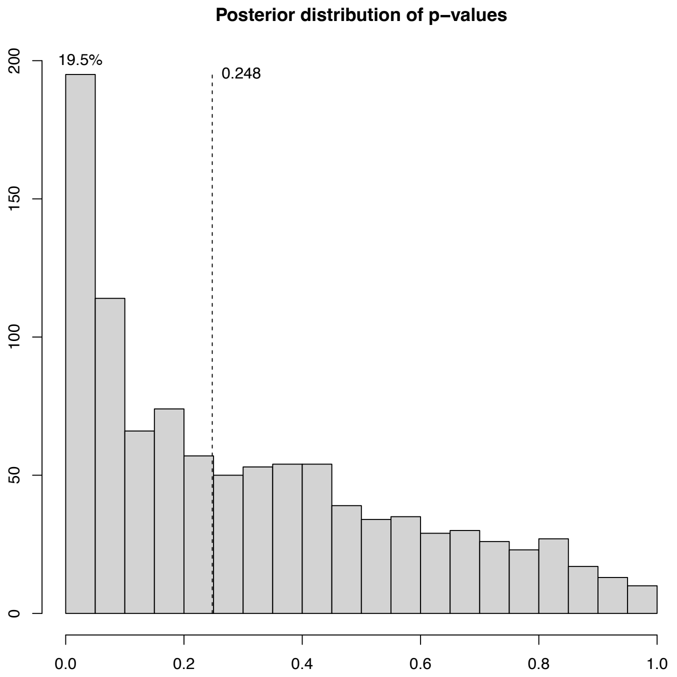


Figure S11: Posterior distribution of residual p-values for the PMEN1 application.

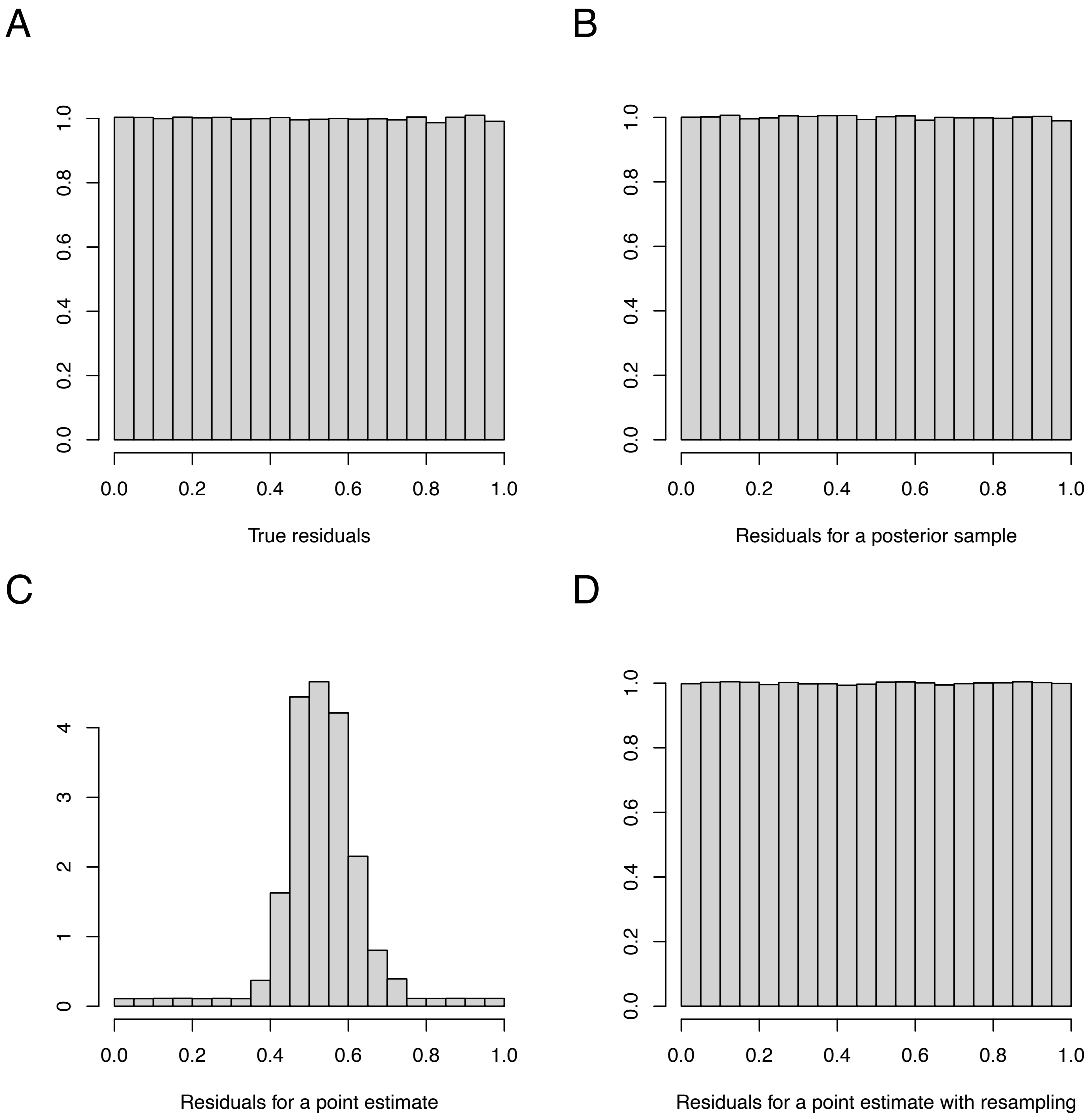


Figure S12: Uniform residuals in the independent and identically distributed case. (A) True residuals. (B) Residuals based on a posterior sample. (C) Residuals based on a point estimate. (D) Residuals based on a point estimate with resampling.
